# SonoMyoNet: A Convolutional Neural Network for Predicting Isometric Force From Highly Sparse Ultrasound Images

**DOI:** 10.1101/2022.06.10.495590

**Authors:** Anne Tryphosa Kamatham, Meena Alzamani, Allison Dockum, Siddhartha Sikdar, Biswarup Mukherjee

## Abstract

Ultrasound imaging or sonomyography has been found to be a robust modality for sensing muscle activity due to its ability to directly image deep-seated muscles while providing superior spatiotemporal specificity compared to surface electromyography-based techniques. Quantifying the morphological changes during muscle activity involves computationally expensive approaches to track muscle anatomical structures or extracting features from B-mode images and A-mode signals. In this paper an offline regression convolutional neural network (CNN) called SonoMyoNet for estimating continuous isometric force from sparse ultrasound scanlines has been presented. SonoMyoNet learns features from a few equispaced scanlines selected from B-mode images and utilizes the learned features to accurately estimate continuous isometric force. The performance of SonoMyoNet was evaluated by varying the number of scanlines to simulate the placement of multiple single element ultrasound transducers in a wearable system. Results showed that SonoMyoNet could accurately predict isometric force with just four scanlines and is immune to speckle noise and shifts in the scanline location. Thus, the proposed network reduces the computational load involved in feature tracking algorithms and estimates muscle force from global features of sparse ultrasound images.

## I. Introduction

Ultrasound is widely used for musculoskeletal imaging and dynamic assessment of muscles. Ultrasound imaging-based sensing of muscle activity, or sonomyography, is recently gaining popularity in prosthetic and rehabilitation applications. Ultrasound imaging allows visualization of muscles along with the anatomical features which play a vital role in muscle contraction during volitional movement. Quantifying changes in muscle morphology during contraction allows estimation of muscle force, and activation level. Estimation of muscle force is essential for understanding biomechanics, for muscle modelling, and for efficient control strategies for prostheses and rehabilitation devices. Estimation of muscle force, joint angles, and joint torques by directly tracking the muscle architectural features in the B-mode ultrasound images has been widely reported. Pennation angle, fascicle length, thickness are the common architectural features that were used to estimate muscle contraction levels of different muscles [2], and joint kinematics of both upper [3], [4] and lower extremities [5], [6]. Hallock et al. quantified the volumetric and spatial deformation of bicep muscles in terms of cross section area, eccentricity of cross section, and thickness to estimate the isometric force [7], [8]. These studies utilized cross correlation tracking algorithms [3], optical flow algorithms [9], [10], [8], [11], beamlet transform-based algorithm [12] etc. to track the architectural features of the muscle. Muscle activity has also been estimated using features from B-mode images. Pixel gradients [13] are few features that were used for identifying muscle activity from B-mode images. A-mode ultrasound-based muscle activity detection is of interest for wearable human machine interfaces. Several studies demonstrated the use of A-mode signals for gesture classification and proportional force estimation. These methods use metrics such as linear fitting coefficients [14], mean and standard deviation [15], [16], [17], waveform length [18] as features to train classifiers or regression models.

On the other hand, convolutional neural networks (CNN) have also been investigated for use in gesture classification as well as continuous joint angle prediction from surface electromyography signals and ultrasound images, due to their ability to learn features directly from image (2D) data without explicit feature extraction. For example, Chen et al. and Ameri et al. have demonstrated surface electromyographybased gesture recongition with CNNs [19], [20]. Ameri et al. used regression-CNNs to implement EMG-based simultaneous wrist motion control which was reported to outperform the support vector machine-based approaches [21], [22]. CNNs with muscle ultrasound images have been used for tracking muscle anatomical structures like cross section area [23], [24]. However, for practical sonomyography systems with only a few ultrasound transducers [25], [15] the performance of a CNN based approach for force prediction is yet to be studied.

Therefore, in this paper, we present a convolutional neural network called SonoMyoNet for quantifying isometric force from sparse ultrasound images of forearm muscle activity. Ultrasound images were acquired from the forearm of healthy subjects during ramped isometric contractions. Equispaced scanlines were extracted from the 2D ultrasound images to create a sparse ultrasound representation of the muscle activity analogous to wearable ultrasound systems with limited transducer count. SonoMyoNet was trained to estimate isometric force exerted by the user with the sparse ultrasound images. The method obviates the need for engineering and extracting features from the ultrasound images. The effect of the number of scanlines and the width of the scanlines on force prediction accuracy have been systematically explored. Practical considerations such as transducer shifts and noise due to speckle have also been evaluated. The following sections describe the the experimental setup, SonoMyoNet network architecture, training and testing methods and results obtained form the study in details.

## II. Methods

### A. Subjects

Five able-bodied subjects with a mean age of 27±3 years with no history of neuromuscular disorders were recruited for the stud. The demographics of the subjects are listed in Table. I. Two subjects (S1 and S2) participated in the experiment twice in two separate sessions. All the participants of the study provided the informed consent before the experiment. The experiment protocols were approved by the Institutional Review Board at George Mason University.

**TABLE I.** Demographics of study participants

| Subject ID | Age | Sex | Dominant arm |
| --- | --- | --- | --- |
| S1* | 30 | Male | Right |
| S2* | 27 | Male | Right |
| S3 | 23 | Male | Right |
| S4 | 24 | Male | Right |
| S5 | 31 | Male | Right |

\* Participated in two sessions at least 4 months apart

### B. Experiment setup

The experimental setup is shown in Fig.1. The subjects were seated comfortably with their forearm placed on a platform and the elbow aligned directly below the shoulder. The forearm was restrained with the help of straps to avoid movement during the experiment. A linear array ultrasound transducer (16HL7, Terason Inc., Burlington, MA USA) was placed on the forearm and secured with a cuff. The ultrasound transducer was transversely placed on the forearm, aligning the field of view to image the major forearm flexor muscle groups and was configured to an imaging depth of 4cm. The ultrasound transducer was connected to a portable ultrasound imaging system (Terason uSmart 3200t, Terason Inc., Burlington MA, USA) and was interfaced to the computer (Intel Core i7-7700HQ, 16GB RAM, 4GB NVIDIA GeForce GTX 1040Ti) using a USB based video capture system (USB2DVI, Epiphan Systems Inc.). The ultrasound images of size 100× 128 were streamed to a custom designed MATLAB (version 2018b, Mathworks Inc., Natick MA, USA) interface in real time at an average frame rate of 22 fps. The isometric grip forces from the hand were measured using a USB-based handheld dynamometer (HD-BTA, Vernier, Beaverton, OR, USA) that was fixed on the platform. The force measurements were acquired at 100 Hz using a custom labVIEW (Version 2020 SP1, National Instruments, Austin, TX, USA) virtual instrument.

**Fig. 1.**
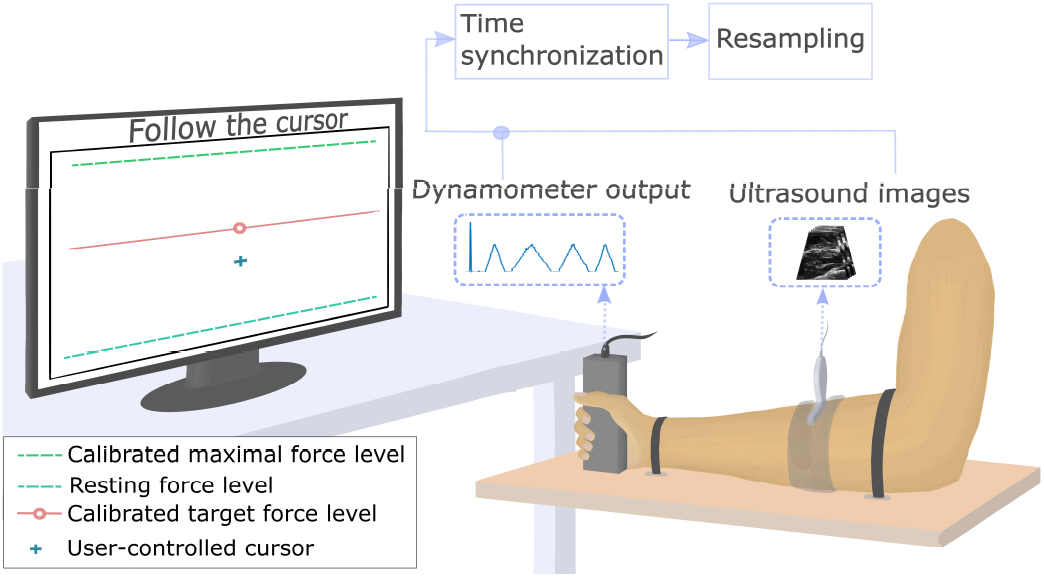
Experimental setup showing the ultrasound probe placed on the subject’s forearm, and the hand dynamometer that measures the isometric grip forces exerted by the subject. The dynamometer force is mapped to the position of the on-screen cursor as the user follows the target. The preprocessing steps involve synchronization and resampling of ultrasound frames and dynamometer force as detailed in [1].

A target tracking task was displayed on a computer screen for the participants. The on-screen task consisted of a target and a user controllable cursor. The dynamometer force readings were mapped to the vertical on-screen position of the user controlled cursor. Subject specific upper and lower bounds for force levels were obtained during an calibration session. Subjects were instructed to perform a maximum voluntary isometric contraction (MVIC) and exert force on the dynamometer for 5 secs followed by rest for 5 secs. The dynamometer force and ultrasound frames corresponding to the MVIC and rest states were recorded during this session. The dynamometer force values corresponding to 80% MVIC and rest were used to normalize the dynamometer readings between ’1’ and ‘0’ respectively. The maximal force was derated to reduce muscle fatigue due to repeated cyclic contractions during the experiment.

During the experiment subjects were instructed to follow the target using the cursor, by modulating the grip force exerted on to the dynamometer. The target cyclically moved between the normalized force levels of ’0’ and ‘1’ at 3 different target velocities. The velocities were chosen such that each ramp cycle (up and down) lasted 20 sec, 40 sec, and 60 sec. Each target velocity was presented thrice with a 10 sec rest phase between each ramp cycle. The ordering of the target velocity was randomized. The dynamometer and the ultrasound frames were simultaneously acquired during the experiment. In order to allow post-hoc synchronization of the dynamometer and ultrasound data streams, subjects were instructed to perform short duration twitches at the start and end of the experiment. Data processing was performed offline as described in the following section.

### C. Ultrasound image and dynamometer data pre-processing

The pre-processing steps for the data were described in detail in [1] and are briefly summarized here. The ultrasound image frames and dynamometer output were time synchronized by aligning the twitch markers at the start and end of the experiment and then resampled to match the ultrasound frame rate . Following the preprocessing steps, sparse ultrasound images were created from the B-mode ultrasound images as shown in Fig. 2. *N* equispaced scanlines with width of *w* were extracted from the B-mode ultrasound images of size 100 × 128, resulting in *N* scanlines of size 100 × *w* . Each scanline, was compressed by computing the mean along the width. The resulting scanlines were concatenated to form a sparse 100 × *N* representation of the B-mode ultrasound images. In our analysis, *N* = 1, 4, 10 with widths *w* = 1.2*mm*, 2.8*mm*, 4.*4mm* The scanlines are analogous to equispaced linear ultrasound transducers placed on the surface of the skin.

**Fig. 2.**
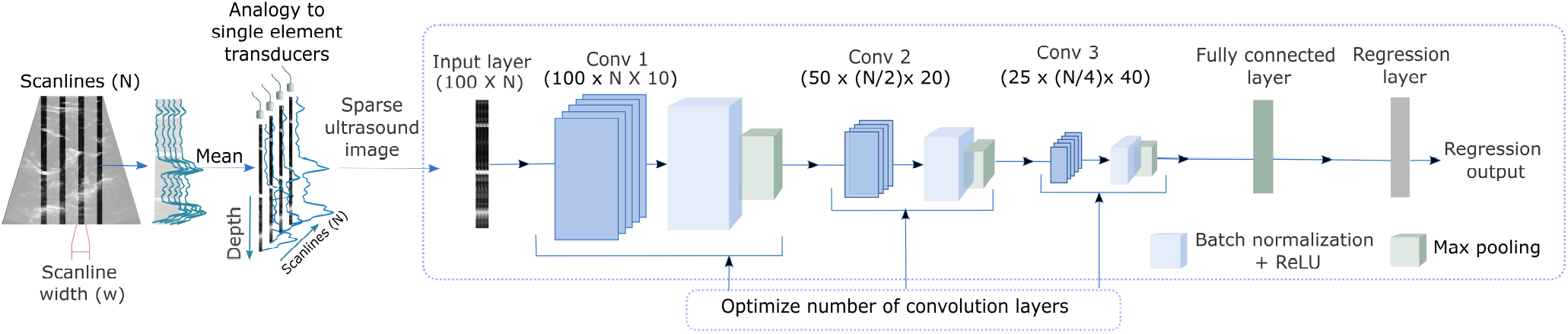
Block diagram demonstrating the sparse image reconstruction from ultrasound images from *N* scanlines of width, *w.* The sparse ultrasound images are fed as inputs to an optimized, subject-specific SonoMyoNet consisting of 4 convolutional layers, followed by a fully connected and regression layer.

### D. Convolutional neural network architecture and optimization

A convolutional neural network (CNN) was designed to predict isometric forces from the sparse ultrasound image data. The CNN architecture used for the analysis is shown in Fig.2. It consists of at least 12 layers. The sparse ultrasound images (100 × N) were first convolved with 10 filters. The output of this convolution layer was batch normalized and ReLU activated. The activation layer output was followed by max pooling. The max pooled features were again convolved using 20 filters and passed through a successive batch normalization, ReLU activation, and max pooling layers. The features were then passed to a third convolution layer with 40 filters, batch normalization, ReLU layers and max pooling layer. The output was fed to a fully connected layer which was followed by a regression layer. For all the convolution layers, a kernel size of 2 × 2 was used with stride ’1’ with ’same’ padding. Stride ’2’ was used for max pooling.

The CNN architecture and training options were optimized for each subject. Sparse ultrasound images and dynamometer force response data were randomly partitioned and 80% of the data was used for training and 20% for validation. The parameters that were optimized were the convolution layer depth, initial learning rate, momentum, and L2 regularization. A Bayes optimization function was used to obtain the optimal parameters by minimising the root mean square error (rmse) of the predicted force.The optimized CNN architecture and training options that were obtained from this process were used in training the CNN with minimum batch size of 150 for 30 epochs.

The experimental data consisting of three trials of ramp up and ramp down cycles performed at three different rates resulted in a total of 9 cycles. Hence, a 9-fold cross validation technique was used to test the prediction capabilities of the optimized network. 8 out of 9 cycles were used for training the network while the remaining cycle was used for isometric force prediction. The process was repeated such that all the cycles were used at least once for force prediction.

### E. Analysis of the factors affecting force prediction accuracy

The effects of various parameters on the force prediction accuracy of the network were analyzed systematically. The following sections explain the rationale for the choice of the parameters.

#### 1) Number of scanlines (N) and scanline width (w)

As previously stated, *N* equispaced scanlines within the field of view of the ultrasound probe were chosen to emulate signals from single element ultrasound transducers. In order to assess the effect of transducer size, a set of *w* scanlines centered around the *N* scanlines were chosen. The average of the intensities of the set of *w* scanlines was computed in order to emulate the resulting A-mode signal received from *N* single element ultrasound transducers. *N* was varied between 1, 4, 10 scanlines while *w* of 1.2mm, 2.8mm, 4.4mm were considered.

#### 2) Transducer shift

Probe and transducer shift during data capture are typical artifacts in ultrasound measurements, particularly in studies involving user movement [25], [26], [27]. The effect of transducer shift was emulated by programmatically shifting the center of the *N* selected scanlines in each frame. A random lateral shift was introduced in all scanlines. The shift was at least one scanline to a maximum shift of six scanlines in either directions The network performance was evaluated by training the network with shift-free ultrasound images and by testing with images corrupted due to transducer shift. The effect of transducer shift was evaluated for different number of transducers and transducer widths.

#### 3) Ultrasound signal to noise ratio

: Speckle is an inherent noise in ultrasound images which is introduced due to presence of sub-wavelength scatterers in the tissue. Multiplicative speckle noise was added to the sparse ultrasound images in the test data set. The speckle noise variance was altered to degrade the peak signal to noise ratio of the sparse ultrasound images from 60 dB to 10 dB. The network was trained with noise-free ultrasound images and tested with the images with speckle noise. The predictive performance of the network was evaluated at each noise level.

### F. Evaluation metrics and statistical analysis

Coefficient of determination (*R*^2^) was used as an evaluation metric for all analysis. The *R*^2^ value is calculated as,

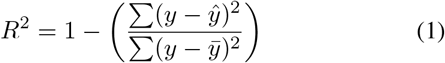

Where, y is the isometric force measured by the dynamometer, *ŷ* is the force predicted by the CNN from sparse ultrasound images in the testing data set, and *ŷ* is the mean value of the measured force values of the testing data set. Non-parametric statistical tests were performed due to small sample size. Friedman’s test was used to compare the *R*^2^ values obtained in different test conditions. Wilcoxon’s signed-rank test was used to compare factors affecting the force prediction. All statistical analysis were performed in IBM SPSS Statistics (Version 28.0, IBM Corp, Armonk,NY, USA).

## III. RESULTS

### 1) Force prediction using CNN

Isometric force was estimated from sparse B-mode ultrasound images using an optimized CNN for all subjects across multiple test conditions. A mean *R*^2^ value of 0.91 ± 0.028, averaged over all scanlines (*N*) and scanline widths (*w*), was obtained using the proposed CNN-based approach for predicting isometric force. Fig.3 shows the time series plot of the measured and predicted forces for one representative subject (S1) for *N* = 1, 4, 10 having a width (*w*) of 4.4mm. The *R*^2^ values obtained from 9-fold cross-validation for *N* = 1, 4, 10 scanlines, each of width 4.4mm, were 0.908±0.05, 0.97±0.01, 0.96 ± 0.017 respectively.

**Fig. 3.**
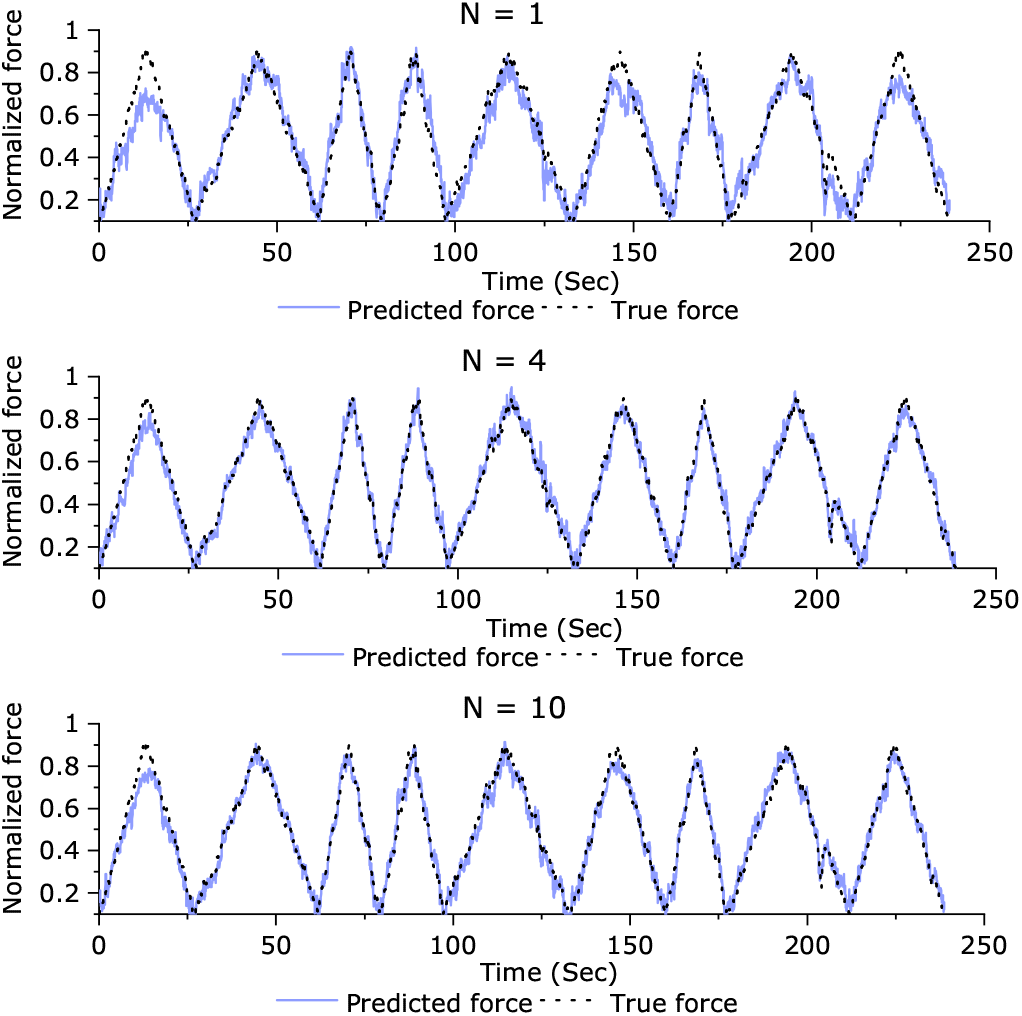
Time series plot showing the measured and predicted forces for one representative subject (S1). A 9-fold cross validation resulted in an *R*^2^ values 0.908±0.05, 0.97±0.01, 0.96 ± 0.017 for 1,4, 10 scanlines having a width of 4.4mm.

### 2) Effect of number of scanlines and scanline width

The effect of number of scanlines (*N*) and channel width (*w*) on force prediction was systematically analysed. Fig.4 shows the *R*^2^ values obtained from force prediction across all subjects when the number of channels and the channel widths were varied. There was a significant improvement in *R*^2^ when the number of channels (*N*) of were increased from 1 to 4 across all channel widths (*p* < 0.001). However, the improvement in *R*^2^ was not significant (*p* > 0.198) when the channels (*N*) were increased from 4 to 10 irrespective of the channel width. Increasing the channel width significantly improved the *R*^2^ value when 1 scanline (*p* = 0.032), 4 scanlines (*p* = 0.004) or 10 scanlines (*p* < 0.001) were employed.

**Fig. 4.**
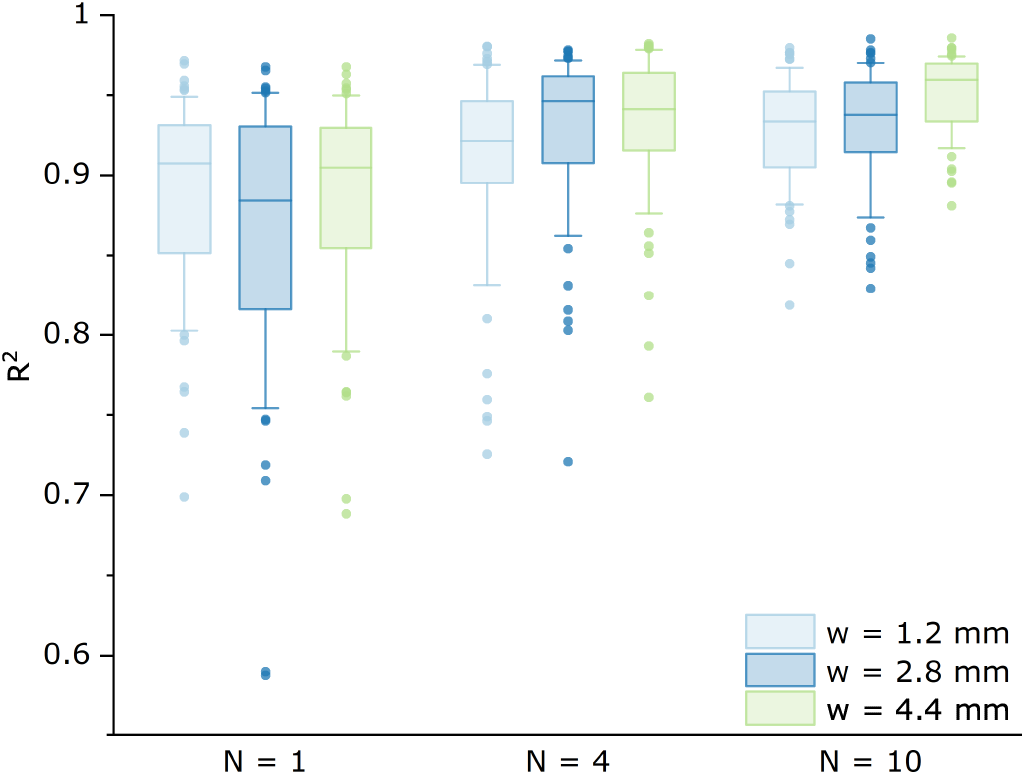
Plot showing the effect of number of scanlines (*N*) and scanline width (*w*) on force prediction performance. *N* = 1, 410 scanlines were considered for analysing the effect of number of scanlines. For each scanline, widths of 0.8mm, 1.4 mm, 3.2 mm were considered.

### 3) Effect of scanline shift

Fig.5 shows the variation in *R*^2^ values due to scanline shift when the number of scanlines (*N*) and the scanline width (*w*) were systematically varied. The equispaced scanlines were randomly shifted from 0.4mm to 2.4mm towards either sides to simulate the effect of random shift in the single element transducer positions. There was a statistically significant effect of scanline shift on the predictive power of the network (*R*^2^) (*p* < 0.001) as seen in Fig.5. In all cases, scanlines with higher widths provided better *R*^2^ than lower widths. However, this difference in performance was significant (*p* < 0.05) for *N* = 1 and 10 scanlines fro shifts up to 1.2mm. For higher shifts,ie., shifts > 1.2mm, having higher scanline width did not provide significantly better performance compared to scanlines with lower width (p > 0.05). It was also observed that scanlines with higher width (i.e. 4.4mm) are immune to transducer shift as seen in Fig.5. For example, for 10 scanlines, shift of 1.2mm resulted in an *R*^2^ of 0.81 ± 0.03 for width of 4.4mm compared to *R*^2^ of 0.65 ± 0.1 for a width of 1.2mm. However at the highest shift of 2.4mm, scanline width did not have significant (*p* = 0.156) effect on the performance of the network. When the scanline width is small (*w* = 1.2mm), increasing the number of scanlines significantly (*p* = 0.043) improved the prediction performance at all shifts.

**Fig. 5.**
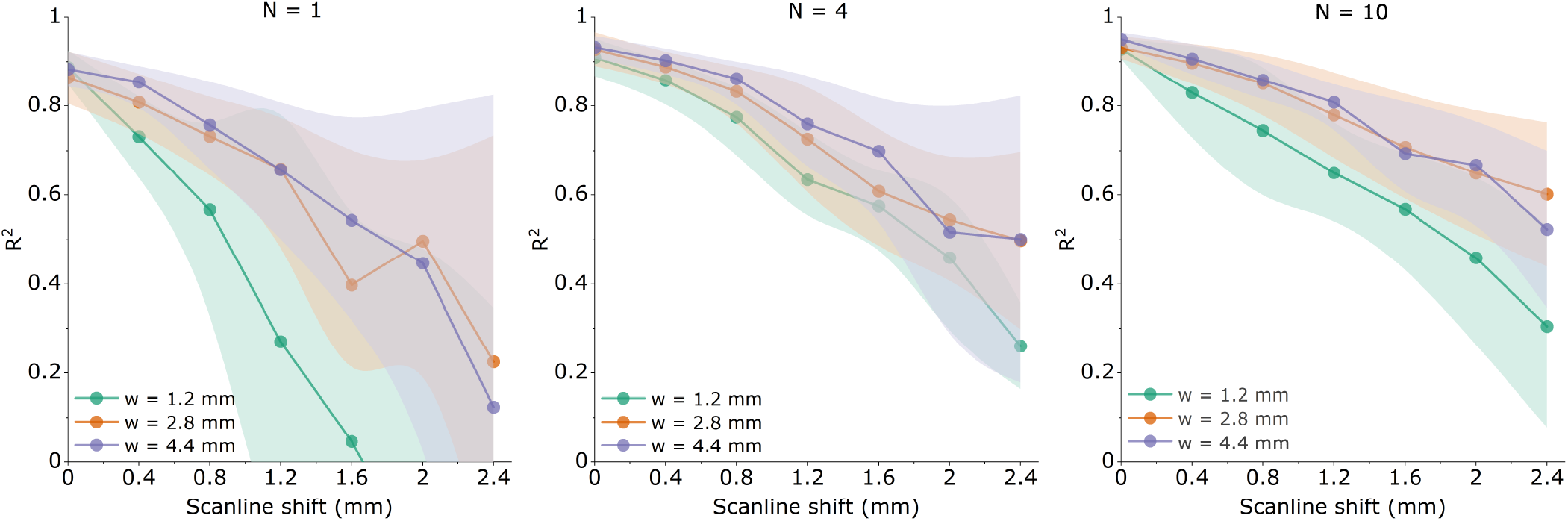
Plot showing the effect of scanline shift on *R*^2^. The SonoMyoNet was trained with images without shift and tested with a test set in which randomized scanlines shifts ranging from 0.4mm to 2.4mm were introduced.

### 4) Effect of speckle noise

Fig.6 shows the effect of speckle noise on force prediction for various combinations of scanlines and widths. There was no significant (*p* > 0.5) effect on *R*^2^ when the PSNR dropped from 62dB to 43dB across all scanlines and widths. However, further detorioration of PSNR by approximately 10dB from 43dB to 33dB resulted in a significant(*p* < 0.5) decrease in *R*^2^.

**Fig. 6.**
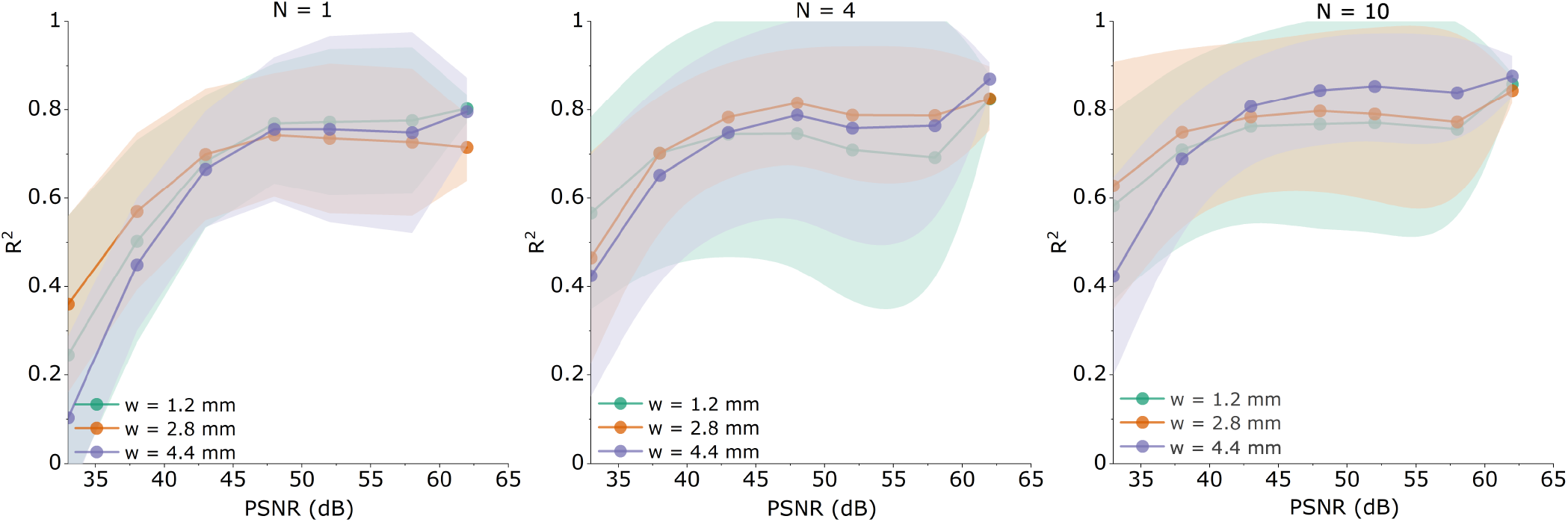
Plot showing the effect of speckle noise on *R*^2^. The SonoMyoNet was trained with speckle noise-free images and tested with images corrupted by speckle noise of different variances that degraded the image PSNR from 63dB to 33dB.

## IV. Discussion

In this paper, a CNN-based network, SonoMyoNet was developed for estimating isometric force from highly sparse B-mode ultrasound images. The performance of SonoMyoNet was evaluated by varying the number of scanlines and scanline width. The effect of common noise and artifacts were also evaluated by introducing shifts in the scanline location and speckle noise conditions. Results demonstrated that the SonoMyoNet could estimate isometric force accurately with an average *R*^2^ value greater than 95% (±0.02) when 10 scanlines of width 4.4mm were considered. This approach estimates isometric forces directly from sparse ultrasound images unlike other methods that require tracking of anatomical structures, or extracting image features to estimate force.

### A. Higher number of scanlines and scanline width provides better force prediction performance

The results demonstrate that increasing the number of scanlines and their width results in better force prediction performance for SonoMyoNet. However, employing more than 4 scanlines does not significantly improve the force prediction performance. Similar results were reported in [1] where, regression models were used to predict isometric force and addition of more scanliens beyond four, did not result in significant performance gains. These results seem to indicate that in a practical system employing single-element ultrasound transducers, the size of the transducers as well as the number of transducers may be critical to accurate force prediction capabilities.

### B. Force prediction accuracy is adversely affected by scanline shift

Practical muscle-computer interfaces employing surface electromyography and ultrasound based systems often suffer from performance degradation due to shifts in the transducer or electrodeposition [22], [28], [29]. Therefore, the performance of SonoMyoNet was evaluated by introducing random shifts in the scanline locations in order to emulate electrode shift. The results show that force prediction is significantly effected by scanline shift. It was observed that a channel shift as small as 2.4mm significantly deteriorated the *R*^2^ value irrespective of number of transducers and transducer width. However, in general, higher scanline count of higher width resulted in higher resilience to shifts.

### C. SonoMyoNet can accurately predict force in the presence of speckle noise

Several studies have evaluated the effect of thermal noise, electrode noise, power line interference etc. in surface electromyography based systems for gesture recognition [30], [31], [32]. Similarly, ultrasound signals are also affected by inherent noise due to speckle. Since speckle is a result of sub-resolution scatterers in tissue, it is critical to evaluate the effect of speckle noise on the performance of the network. It was observed that the *R*^2^ values obtained when the force was predicted using images with speckle noise were lower than those predicted using images without speckle noise. However, SonoMyNet was found to be resilient to speckle noise of approximately 20dB with no significant degradation in performance across. Therefore, the proposed method could predict force with acceptable *R*^2^ values even from images having a PSNR of 38dB.

### D. Limitations

SonoMyoNet provides a robust, noise-tolerant and subject-specific method for estimation of continuous isometric force from sparse reconstruction of ultrasound images. While this method provides several key advantages, the CNN architecture has certain limitations. The network requires several iterations of ramped isometric contractions to generate training data and ensure convergence. Repeated isometric contractions may lead to muscle fatigue, thus leading to model inaccuracies. In this work the effect of fatigue was minimized by derating the maximal applied force during training. Further, this work tests the performance of the network for offline estimation of isometric force. It has been shown that online performance in classification and regression tasks for surface electromyography based human-machine systems shows significant degradation compared to offline results. It is expected that further optimization may be required for SonoMyoNet architecture for online force estimation applications.

## V. Conclusion

SonoMyoNet provides a robust method for estimation of isometric forces from highly sparse representations of ultrasound images. The network is demonstrated to possess a high degree of immunity to common noise sources such as speckle noise and electrode shifts. This work also establishes design parameters for a practical wearable ultrasound based muscle activity measurement system employing a CNN-based approach. This work paves the way for isometric force estimation from a collection of A-mode ultrasound transducer without the need for feature extraction.

